# *Drosophila* R8 photoreceptor cell subtype specification requires *Notch* and *hibris*

**DOI:** 10.1101/398222

**Authors:** Hong Tan, Ruth E. Fulton, Wen-Hai Chou, Denise A. Birkholz, Meridee P. Mannino, David M. Yamaguchi, Steven G. Britt

## Abstract

Cell differentiation and cell fate determination in sensory systems are essential for stimulus discrimination and coding of environmental stimuli. Color vision is based on the differential color sensitivity of retinal photoreceptors, however the developmental programs that control photoreceptor cell differentiation and specify color sensitivity are poorly understood. In *Drosophila melanogaster,* there is evidence that the color sensitivity of different photoreceptors in the compound eye is regulated by inductive signals between cells, but the exact nature of these signals and how they are propagated remains unknown. We conducted a genetic screen to identify additional regulators of this process and identified a novel mutation in the *hibris* gene. *hibris* encodes an *irre* cell recognition module protein (IRM). These immunoglobulin super family cell adhesion molecules include human neph and nephrin (NPHS1). *hibris* is expressed dynamically in the developing *Drosophila melanogaster* eye and loss-of-function mutations give rise to a diverse range of mutant phenotypes including disruption of the specification of R8 photoreceptors cell diversity. The specification of blue or green sensitivity in R8 cells is also dependent upon *Notch* signaling. We demonstrate that *hibris* is required within the retina, non-cell autonomously for these effects, suggesting an additional layer of complexity in the signaling process that produces paired expression of opsin genes in adjacent R7 and R8 photoreceptor cells.

**Author Summary:** As humans, our ability to distinguish different colors is dependent upon the presence of three different types of cone cell neurons in the retina of the eye. The cone cells express blue, green or red absorbing visual pigments that detect and discriminate between these colors. The principle of color discrimination by neurons “tuned” to different colors is an evolutionarily conserved specialization that occurs in many different animals. This specialization requires 1) visual pigments that detect different colors and 2) a developmental program that regulates the expression of these pigments in different types of cells. In this study we discovered that the fruit fly (*Drosophila melanogaster*) gene *hibris* is required for the developmental program that produces blue sensitive neurons in the fly retina. When we over-expressed *hibris* throughout the developing retina, extra blue sensitive cells were produced. These results demonstrate that if there is not enough *hibris,* too few blue sensitive cells form, but if there is too much *hibris*, too many blue sensitive cells form. Finally, we discovered that the *hibris* gene does not act in color sensitive neurons of the retina themselves. This surprising discovery suggests that *hibris* may influence development of the retina in a completely new and different way.

## Introduction

Color vision in humans and most other organisms is dependent upon the expression of spectrally distinct visual pigments (opsins) in different photoreceptor cells [1-3]. The organization of photoreceptor cells within the retinal mosaic reflects a variety of different developmental mechanisms, including regional specialization, stochastic, and precise cell-cell adjacency [4]. *D. melanogaster* is capable of color vision and is a useful experimental system for examining the developmental programs that produce photoreceptor cells having different color sensitivities [5-12]. The compound eye consists of ∼800 ommatidia, each containing eight rhabdomeric photoreceptor cells (R cells). The central R7 and R8 photoreceptor cells mediate polarization sensitivity and color vision [13, 14]. As shown in **Fig 1**, the majority of ommatidia contain matched pairs of R7 and R8 cells expressing specific rhodopsin (Rh) visual pigments, either *Rhodopsin 3* (*Rh3*, FBgn0003249) and *Rhodopsin 5* (*Rh5*, FBgn0014019) (tandem magenta-blue cylinders), or *Rhodopsin 4* (*Rh4*, FBgn0003250) and *Rhodopsin 6* (*Rh6*, FBgn0019940) (tandem yellow-green cylinders).

**Fig 1.**
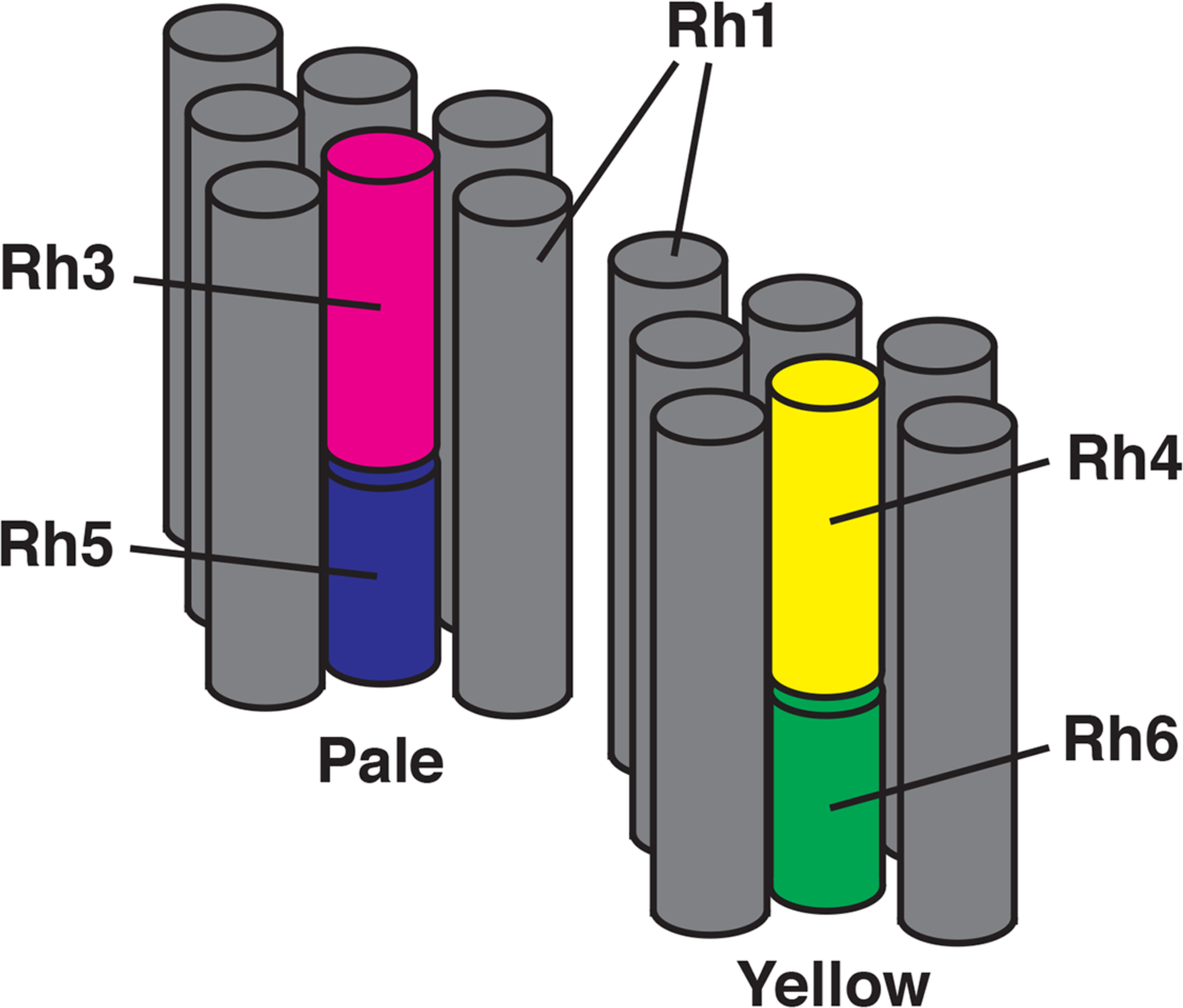
Diagram of photoreceptor cell organization and opsin gene expression. Two ommatidia are shown consisting of gray cylinders corresponding to the rhabdomeres of the R1-6 photoreceptor cells. These surround the central rhabdomeres of the R7 and R8 cells. Expression of opsin genes within the R7 cells (*Rh3* in magenta or *Rh4* in yellow) is paired with opsin gene expression in the adjacent R8 cell (*Rh5* in blue or *Rh6* in green) in pale and yellow ommatidia, respectively.

These two main ommatidial subtypes were initially identified based on pale or yellow fluorescence when illuminated with blue light [15, 16], with pale (pR7/pR8) expressing *Rh3/Rh5*, while yellow (yR7/yR8) cell pairs express *Rh4/Rh6* (**Fig 1**) [10, 11, 17]. This paired expression of opsin genes in adjacent R7 and R8 cells within an individual ommatidium is thought to result from a series of developmental steps. First, a subset of R7 cells stochastically and cell autonomously express *spineless* (*ss,* FBgn0003513) which represses *Rh3* and induces *Rh4* expression [18]. In pR7 cells that stochastically fail to express *ss* and do express *Rh3*, a signal is initiated that induces the expression of *Rh5* in adjacent pR8 cells. Extensive studies have identified the genes *warts (wts,* FBgn0011739), *melted (melt,* FBgn0023001), members of the *hippo (hpo,* FBgn0261456) pathway, along with the TGFβ superfamily receptors *baboon* (*babo,* FBgn0011300) and *thick vein* (*tkv,* FBgn0003726), and their respective ligands as components of the inductive signal from pR7 that drives the expression of *Rh5* in pR8 [12, 19-21]. In the absence of a signal from yR7, the default yR8 fate and expression of *Rh6* occurs. In addition, we have found that the *Epidermal growth factor receptor (Egfr,* FBgn0003731) and *rhomboid (rho,* FBgn0004635) are also required for this process [22, 23].

Here we undertook a genetic screen to identify additional genes required for this process and show that *hibris* (*hbs,* FBgn0029082), an *irre* Cell Recognition Molecule (IRM) [24], NPHS1 (nephrin, *Homo sapiens,* HGNC:9801) related member of the Immunoglobulin Super Family (IgSF), as well as *Notch* (*N,* FBgn0004647) are required for the establishment of paired opsin expression in adjacent R7 and R8 photoreceptor cells. Interestingly, we found that *hbs* is required non-cell autonomously for this process, suggesting the involvement of additional interactions between R7, R8 and neighboring cells.

## Results

### Isolation and characterization of the *a69* mutant

To identify genes required for the induction of *Rh5* expression in R8 photoreceptors, we screened a collection of approximately 150 homozygous viable eye-expressing enhancer trap lines carrying insertions of the *P{etau-lacZ}* transposon (FBtp0001352) [25]. This was based on the rationale that genes required for the induction of *Rh5* expression would be expressed in the eye, the *P{etau-lacZ}* transposon has been especially useful in studies of the nervous system, and insertion of this element into loci of interest would provide a convenient means to identify the affected genes [25]. The percentage of *Rh5*-expressing R8 cells was determined by labeling dissociated ommatidia with antibodies against *Rh5* and *Rh6*. Several mutants with abnormal percentages of *Rh5*-expressing R8 cells were noted and *a69* (FBgn0026612), with the lowest percentage of *Rh5* (9%) was further characterized. Immunostaining of both dissociated ommatidia and tissue sections showed that in the *a69* enhancer-trap line, *Rh5*- expressing R8 cells are reduced and most R8 cells have assumed the default fate and express *Rh6* (**Fig 2A-E**, **Table 1**). Since mutants lacking R7 cells or having a reduced number of Rh3 expressing R7 cells would also show diminished *Rh5* expression, we examined the expression of the opsins expressed in the R7 cells and found that the percentage of *Rh3* expressing R7 cells was similar to *white*^*1118*^ (*w*^*1118*^, RRID:BDSC_3605) control flies (41.9%, **Table 1**).However, there was a dramatic mispairing between *Rh3* expressing R7 cells adjacent to *Rh6* expressing R8 cells (**Fig 2C,F**, **Table 1**) compared to *cinnabar*^*1*^ *brown*^*1*^ controls (*cn*^*1*^ *bw*^*1*^, RRID:BDSC_264) consistent with the idea that the *a69* enhancer trap line carries a mutation in a gene required for the induction of *Rh5* expression in R8 cells.

**Table 1.** Opsin Expression in Different Genetic Backgrounds.

| Genotype | R8 cells expressing *Rh5*<br>% (n) | R7 cells expressing *Rh3*<br>% (n) | Mis-pairing<br>% (n) | Figure |
| --- | --- | --- | --- | --- |
| *w<sup>1118</sup>* | 29 (214) | 47 (362) | *Rh4/Rh5* 0 (424) | 2A, B, C |
| *a69* | 9 (335)<br>SDF *w<sup>1118</sup>*, $p = 1.9 \times 10^{-9}$ | 42 (241) | *Rh3/Rh6* 25 (253)<br>SDF *cn<sup>1</sup> bw<sup>1</sup>*, $p = 1.2 \times 10^{-8}$<br>*Rh4/Rh5* 0 (315) | 2D, E, F |
| *cn<sup>1</sup> bw<sup>1</sup>* | ND | ND | *Rh3/Rh6* 6 (240) |  |

Statistical comparisons of strains were carried out as described in the Methods; n= the number of ommatidia counted. Unless indicated, the observed percentages were not significantly different from *w<sup>1118</sup>*. Strains compared to another control are indicated. Abbreviations are as follows: Significantly Different From (SDF) the strain indicated, at the $p$ value shown by a two tailed test; Not Determined (ND); Not Applicable (NA).

**Fig 2.**
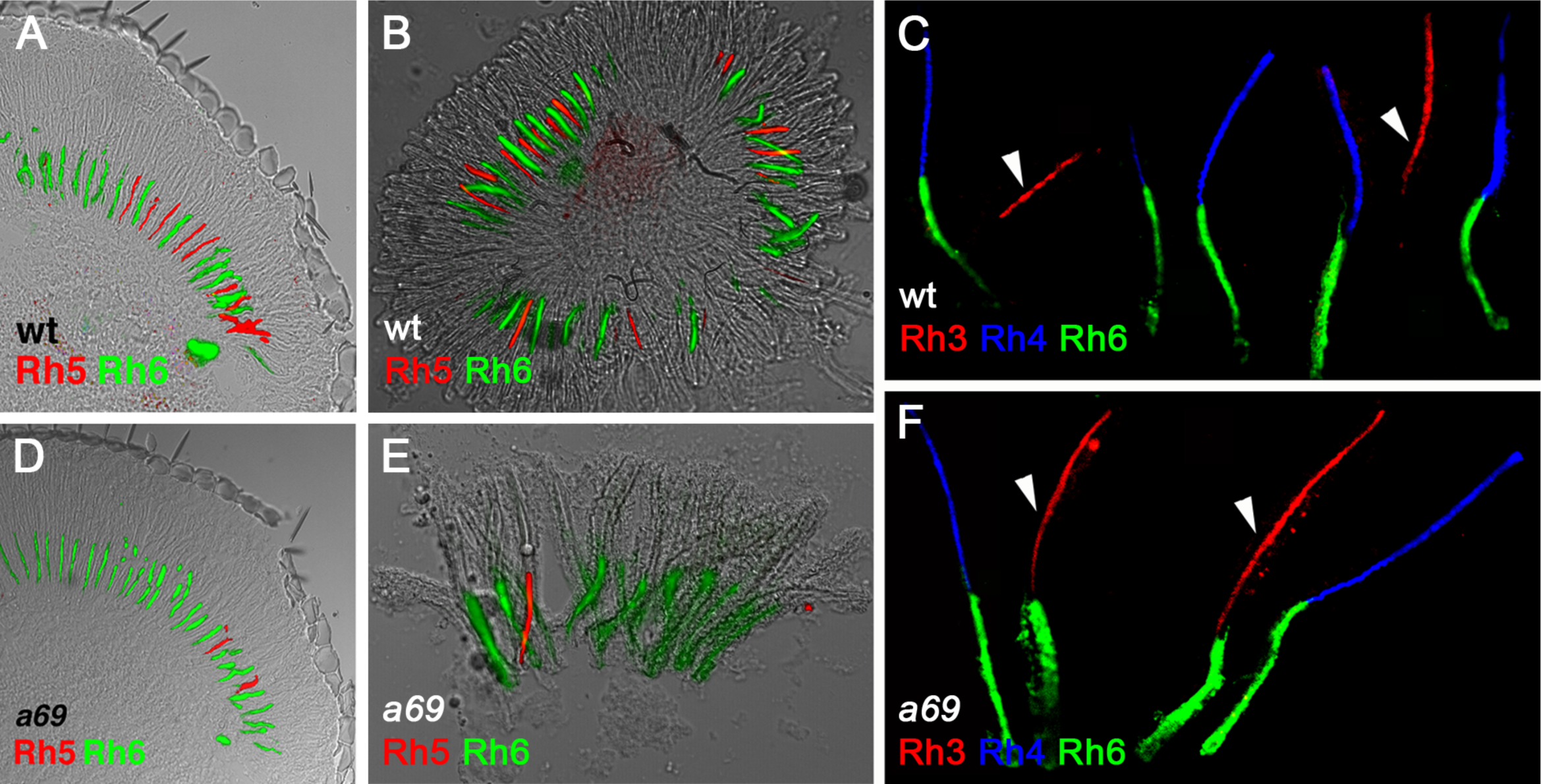
*a69* mutants have a defect in Rh5 and Rh6 expression in R8 photoreceptor cells. White eyed wild type flies (*w*^*1118*^) express *Rh5* and Rh6 in a proportion of approximately 1:2, this is shown in a longitudinal section of the retina (A) as well as in dissociated ommatidia (B). (C) *w*^*1118*^ flies express Rh4 and Rh6 in a paired fashion. The arrowheads indicate Rh3 expressing R7 cells that do not normally pair with Rh6 expressing R8 cells. *w*^*1118*^; *P{etau-lacZ}a69* mutants show a disruption in Rh5 expression, with a substantial decrease in Rh5 expression shown in both section (D) and dissociated ommatidia (E) as well as prominent mispairing between Rh3 expressing R7 cells and Rh6 expressing R8 cells in the same ommatidia (arrowheads).

To isolate the gene responsible for the *a69* phenotype, the location of the P-element insertion in *a69* was determined and found to map to the right arm of the second chromosome at position 60E (data not shown). To determine whether the P-element in *a69* is the cause of the phenotype, P-element excision lines were generated and analyzed. Thirty-five homozygous strains of these excision chromosomes were analyzed by staining dissociated ommatidia with antibodies against *Rh5* and *Rh6*, and all of them (100%) were found to have a low *Rh5* percentage, similar to that of *a69* (data not shown). Only 1% of excision strains would be expected to retain the mutant phenotype as a result of imprecise excision, thus our inability to revert the mutant phenotype is consistent with the *a69* P-element not being responsible for the mutation [26]. Furthermore, mapping via recombination analysis revealed that the *a69* mutation is localized to the interval between the *purple* (*pr,* FBgn0003141) and *curved* (*c,* FBgn0000245) genes in the middle the second chromosome (**Fig 3, S 1 Table**), far away from the P-element insertion site in *a69.* From this we conclude that the *a69* mutation is not associated with the insertion of the P-element. Thirty-three deficiency lines located in the region between *pr* and *c* were tested for *a69* complementation (**Fig 4, S 2 Table**). These analyses narrowed the location of the *a69* mutation to 51C3-51D1 (**Fig 4**). The lower portion of **Fig 4** shows a diagram of this genomic region, spanning ∼300 Kb and encompassing 25 known protein coding genes.

**Fig 3.**
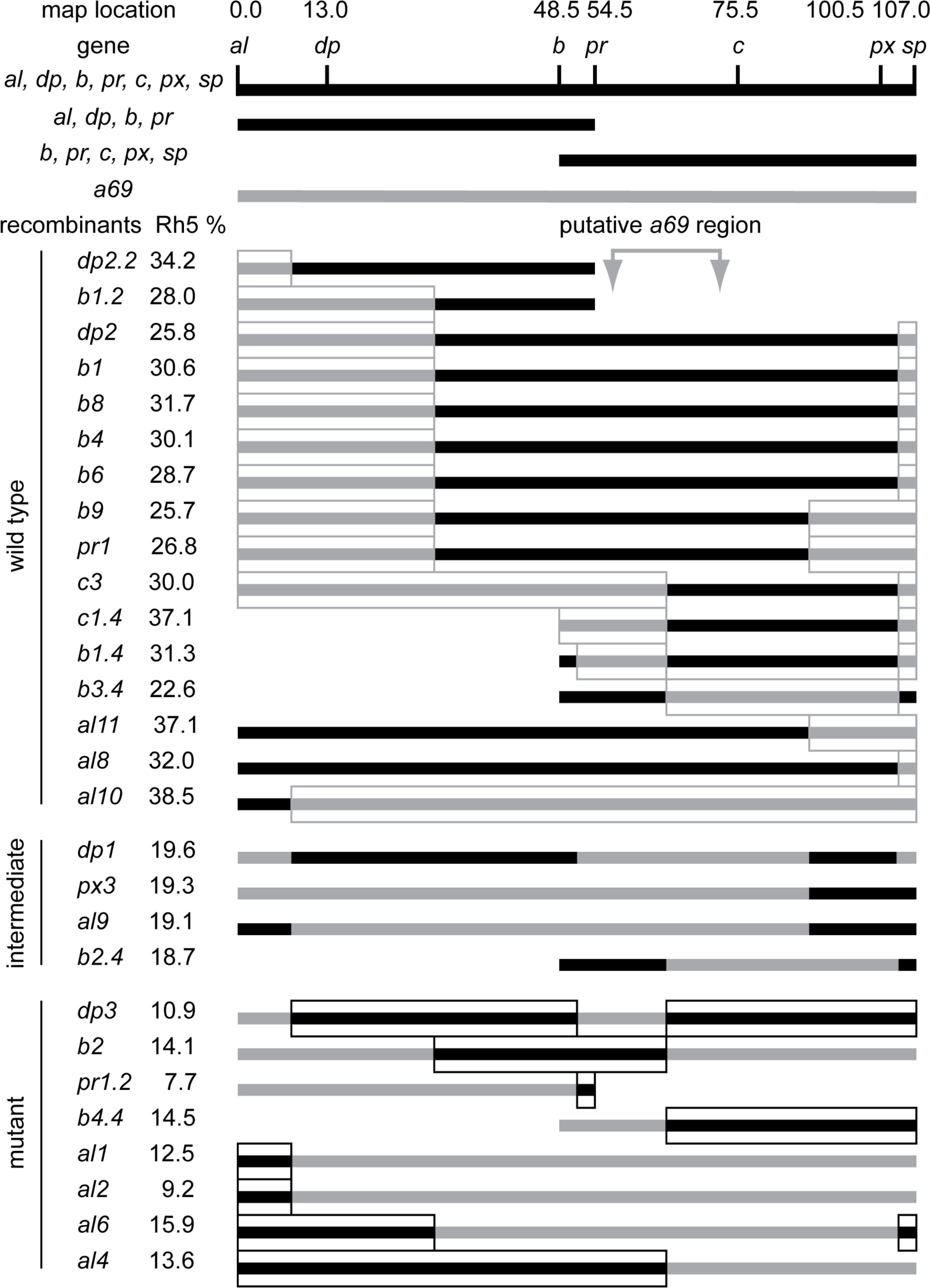
Recombination mapping of *a69* to the second chromosome between *pr* and *c*. Three multiply marked chromosomes (*al*^*1*^ *dpy*^*ov1*^ *b*^*1*^ *pr*^*1*^ *c*^*1*^ *px*^*1*^ *sp*^*1*^, *al*^*1*^ *dpy*^*ov1*^ *b*^*1*^ *pr*^*1*^, and *b*^*1*^ *pr*^*1*^ *c*^*1*^ *px*^*1*^ *sp*^*1*^) were recombined with the *w*^*1118*^; *P{etau-lacZ}a69* mutant. After marker identification, recombinant strains were back crossed to the *a69* mutant and scored for the percentage of Rh5 expression. The regions of the recombinant chromosomes assumed to be derived from the *a69* parental mutant strain are indicated in gray, while the regions assumed to be derived from the multiple marked (wild-type) chromosomes are black. Sixteen recombinant strains were phenotypically wild-type and complemented *a69*. Four recombinant strains were intermediate and eight strains were mutant and failed to complement *a69*. The four intermediate strains and one wild type strain, *al10,* differed from the expected phenotypes and may have resulted from multiple recombination events or exposure of cryptic modifier loci. See **S1 Table**. Complementation of *a69* Recombinant Strains.

**Fig 4.**
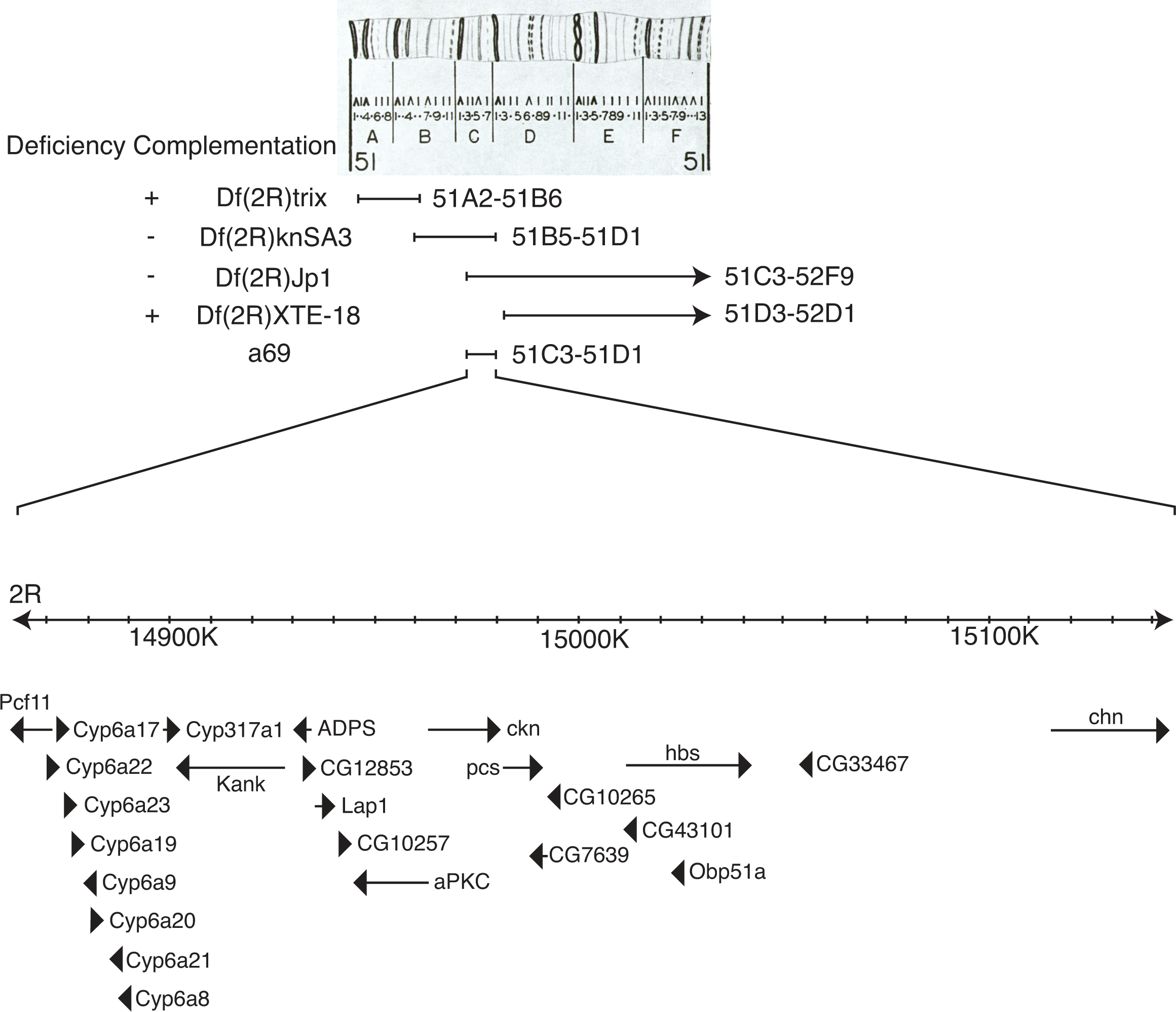
Cytogenetic Map, Molecular Map and Deficiency Complementation of *a69*. The top panel shows the cytogenetic map of the 51 region of chromosome 2R [74], used with permission. Diagramed below are the deleted regions of deficiency strains tested, the corresponding molecular map and identified protein coding genes in the region. Arrows or arrowheads indicate the orientation of gene transcription and arrow or arrowhead length corresponds to gene length at the scale indicated (K, kilobase). Data obtained from Flybase version FB2018_01 [64].

To identify the gene specifically in the *a69* mutation, we took two approaches. First, a subset of genes were examined for alterations in expression in the *a69* mutant, and second, a large series of complementation studies were performed with alleles of known mutants in the region. cDNAs from 5 genes in the region were obtained and *in-situ* hybridization of third instar larval eye imaginal discs was performed on *cn*^*1*^ *bw*^*1*^ (wild-type) and *a69* mutants. In each case the expression pattern of the gene was not substantially disrupted in *a69* mutants, suggesting that the phenotype is not due to the disruption of patterned mRNA expression of these genes in the 3rd instar eye-antennal disc. (**Fig 5**). *hibris* (*hbs*) was expressed strongly in the morphogenetic furrow and maintained weakly posteriorly, consistent with a previous report [27]. It was also expressed in the ocellar region and in the developing antenna. *parcas* (*pcs,* FBgn0033988) was expressed strongly in the morphogenetic furrow and in the antenna. *CG10265* (FBgn0033990) did not appear to be expressed in either the eye or antennal regions. *CG7639* (FBgn0033989) appeared to be weakly expressed in the region anterior to the morphogenetic furrow. *caskin* (*ckn,* FBgn0033987) was expressed anterior to the furrow and in the antenna.

**Fig 5.**
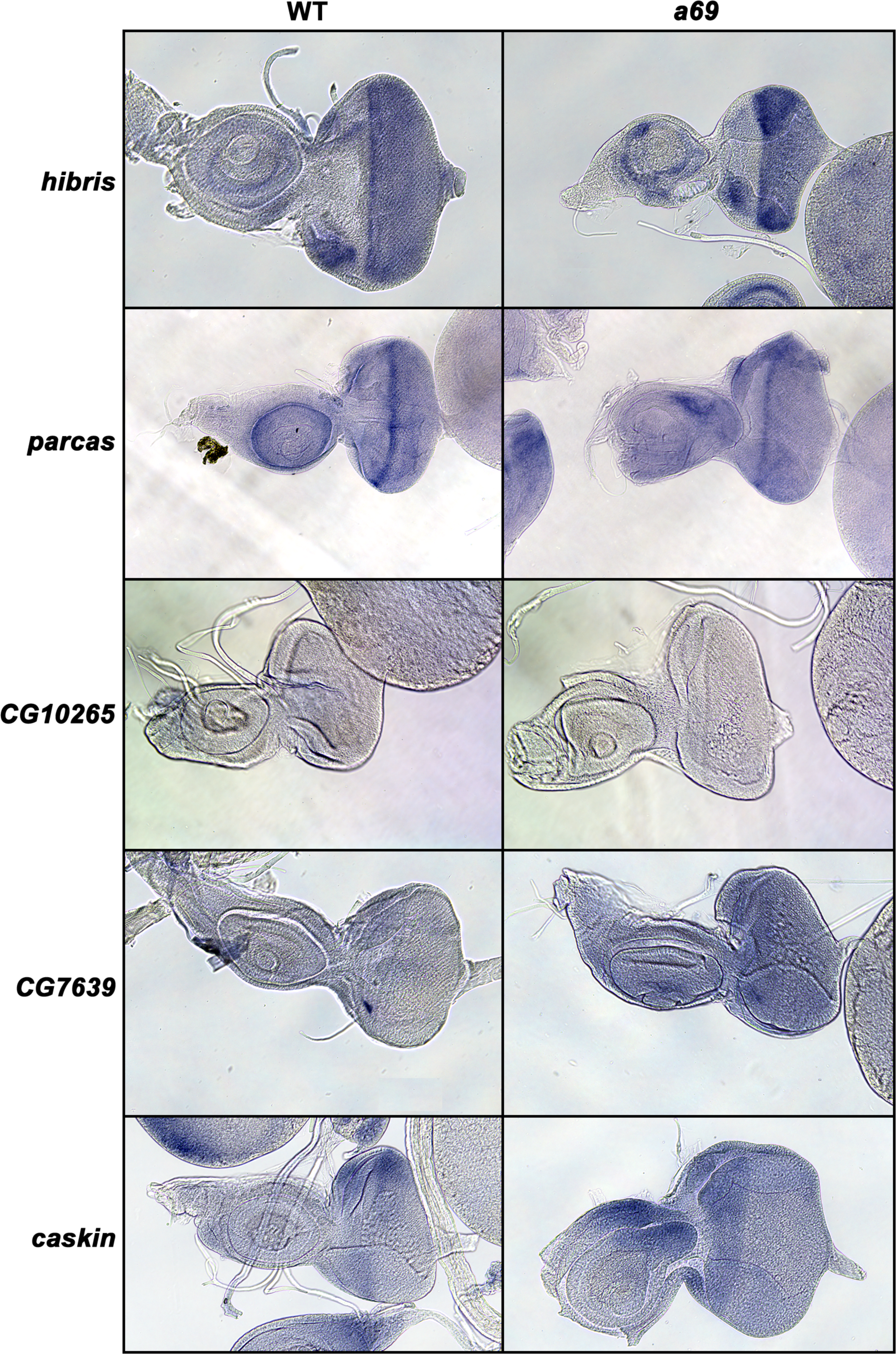
*In situ* Hybridization of *a69* Candidate Genes. The figures shows *in situ* hybridization of biotinylated reverse strand probes prepared from *hibris, parcas, CG10265, CG7639,* and *caskin* cDNA clones (rows) against wild type (*cn*^*1*^ *bw*^*1*^) (left column) or *a69* mutant (right column) eye-antennal imaginal discs.

We characterized *Rh5* and *Rh6* expression in animals heterozygous for *a69* and alleles of *Additional sex combs* (*Asx,* FBgn0261823), *atypical protein kinase C* (*aPKC,* FBgn0261854), *bocce* (*boc,* FBgn0011203), *charlatan* (*chn,* FBgn0015371), *Enhancer of GMR-sina 2-1* (*ES2-1,* FBgn0024358), *Hexokinase C* (*Hex-C,* FBgn0001187), *knot* (*kn,* FBgn0001319), *Regulatory particle non-ATPase 6* (*Rpn6,* FBgn0028689), *safranin* (*sf,* FBgn0003367), *Protein 1 of cleavage and polyadenylation factor 1* (*Pcf11,* FBgn0264962), *scab* (*scb,* FBgn0003326), and transposon insertions P{A26O9}1 (FBti0001751) and P{lacW}B6-2-25 (FBti0005748). All of these mutations complemented *a69* (data not shown).

We obtained the following alleles of *hbs*: *hbs*^*361*^(FBal0130217), *hbs*^*459*^, (FBal0130216), *hbs*^*1130*^(obtained from M. Baylies) and *hbs*^*2593*^ (FBal0130218). With one exception, all of these alleles fail to complement *a69*, **Table 2**. Furthermore, *hbs*^*361*^ homozygotes and heteroallelic combinations of these alleles all show a substantial decrease in the proportion of *Rh5* expression in R8 photoreceptor cells. With four exceptions, viable combinations of these alleles over deficiencies in the region show the same complementation pattern as the *a69* mutant, **S3 Table**.

**Table 2.** **Complementation crosses of *a69, hbs* alleles and *cn bw* control.**

| Genotype of Strains Crossed | *hbs*<sup>361</sup> | *hbs*<sup>459</sup> | *hbs*<sup>1130</sup> | *hbs*<sup>2593</sup> | *cn*<sup>1</sup> *bw*<sup>1</sup> |
| --- | --- | --- | --- | --- | --- |
| *a69* | 5.0% (337) | 22.9% (1164)<br>$p = 1.7 \times 10^{-4}$ | 10.4% (201) | 1.5% (455) | 25.7% (152)<br>$p = 6.4 \times 10^{-4}$ |
| *hbs*<sup>361</sup> | 16.6% (404) | 3.3% (456) | 1.4% (358) | 2.7% (414) | 29.1% (320)<br>$p = 1.4 \times 10^{-6}$ |
| *hbs*<sup>459</sup> | | | 3.9% (799) | 2.5% (651) | 33.3% (699)<br>$p = 1.3 \times 10^{-10}$ |
| *hbs*<sup>1130</sup> | | | | 1.2% (326) | 26.8% (503)<br>$p = 5.2 \times 10^{-6}$ |
| *hbs*<sup>2593</sup> | | | | | 30.7% (703)<br>$p = 8.5 \times 10^{-9}$ |

Statistical comparisons of strains were carried out as described in the Methods. Values shown are percentage of R8 cells expressing Rh5 (number of ommatidia counted). The crossed alleles fail to complement *a69* and each other (shaded gray). Complementation in this table (unshaded) is an *Rh5*% significantly greater than *a69* homozygotes (12.7% (267)) by a one tailed test at the *p* value shown.

Exon sequencing of the *hbs* gene failed to identify unique polymorphisms in the *a69* mutant that were absent in phenotypically wild type control strains (data not shown). Nonetheless, given that the gene spans over 30 Kb including 24 Kb in the first intron, it seems likely that a mutation within a regulatory region of the gene may be responsible for the hypomorphic *a69* phenotype and the complex complementation pattern found with the *hbs*^*459*^ allele. Thus, we believe the complementation data is fully consistent with *a69* being a *hbs* allele, *hbs*^*a69*^.

### *hibris* is expressed in the developing third instar eye imaginal disc

Consistent with previous studies [28], we find that *hbs* is expressed in the developing third instar eye imaginal disc in preclusters of photoreceptor cells emerging from the morphogenetic furrow and ultimately in all photoreceptor cells, **Fig 6**. *hbs* is expressed coordinately with early *senseless* (*sens*, FBgn0002573) expression in R8 just posterior to the morphogenetic furrow and this is followed by *prospero* (*pros,* FBgn0004595) expression in R7 cells 6-8 rows posterior and cone cells.

**Fig 6.**
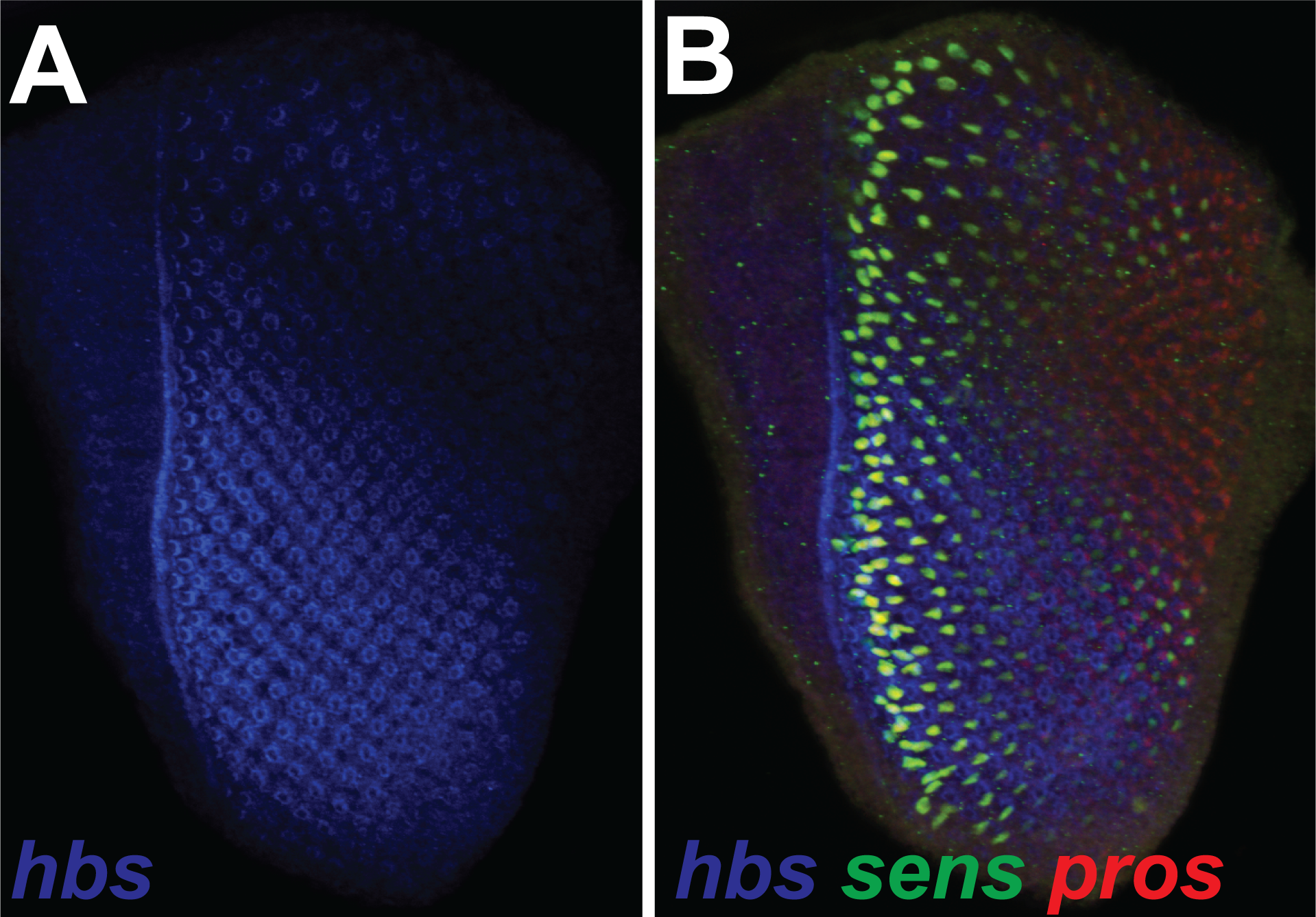
*hibris* Expression in the Third Instar Larval Eye Imaginal Disc. The figure shows a confocal microscopy flattened Z-stack series of *hibris* (*hbs*) immunolabeling (A) and triple labeling of the same wild type (*cn*^*1*^ *bw*^*1*^) specimen with antibodies against *hbs*, *senseless* (*sens*) and *prospero* (*pros*) (B). The morphogenetic furrow has moved from right (posterior) to left (anterior).

### *hibris* is required in the retina for R7 and R8 cell differentiation

To examine the function of *hbs* in Rh5 and Rh6 expression in R7 and R8 photoreceptor cell patterning, we examined an additional allele of *hbs* in mosaic flies. We used the *ey-FLP* driver to generate homozygous mutant clones in the retina and optic lobes of animals that were heterozygous for *hbs*^*1130*^. We used a cell autonomous lethal to generate large homozygous mutant clones and eliminate homozygous wildtype tissue, as described [29]. **Fig 7** shows a small heterozygous clone with a single *Rh5* expressing R8 cell in an otherwise homozygous mutant retina where *Rh3* expressing R7 cells are mispaired with *Rh6* expressing R8 cells, demonstrating a phenotype identical to *a69*.

**Fig 7.**
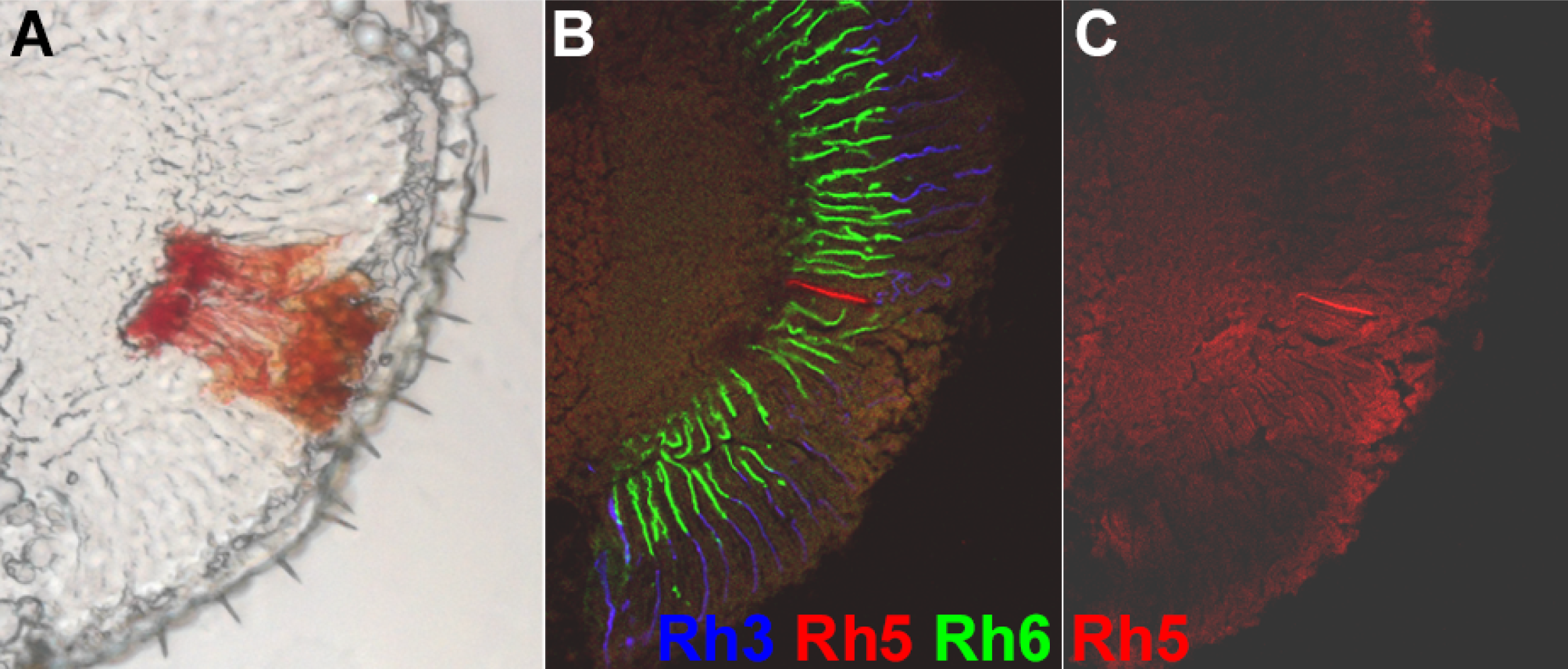
Opsin Expression in *hbs*^*1130*^Mosaic Mutant Retina. Large FLP-FRT retinal clones were generated in the eye and optic lobes with *ey-FLP*. Panel A shows a bright field microscopy image of a single heterozygous clone (red tissue marked with *w*^*+*^) within an otherwise homozygous *hbs*^*1130*^ mutant retina (white tissue). B and C show one R8 photoreceptor cell expressing *Rh5* in this heterozygous clone, whereas elsewhere in the retina, only *Rh6* is expressed in R8 photoreceptor cells. Frequent mispairing of *Rh3* expressing R7 cells and *Rh6* expressing R8 cells is also shown. *Rh3* (blue), *Rh5* (red) and *Rh6* (green) expression were detected by confocal microscopy with directly labeled monoclonal antibodies as described in **Materials and Methods**.

To further refine the spatial requirement for *hbs* in R7 and R8 photoreceptor cell differentiation and opsin gene expression we compared mutant clones of *hbs*^*66*^ (FBal0239852) [30] generated using *ey-FLP* and *ey3.5-FLP* [31*]. ey3.5-FLP* is a modified form of *ey-FLP* that efficiently induces clone formation in the third instar larval eye imaginal disc, but not in the lamina or medulla. **Fig 6A** shows loss of *hbs* in the retina and optic lobe leads to a dramatic decrease in *Rh5* expression and mispairing of *Rh3* and *Rh6* in adjacent R7 and R8 cells of individual ommatidia, consistent with the results obtained with *hbs*^*1130*^, **Fig 5**. This is in contrast to *Rh3*, *Rh5* and *Rh6* expression in a similarly FRT recombined clone of a wild type chromosome. By comparison, retina specific clones generated with *ey3.5-FLP* [31] also show a loss of *Rh5* expression and mispairing of *Rh3* and *Rh6*. These results indicate that *hbs* is required in the retina for normal R7 and R8 photoreceptor cell differentiation and opsin gene expression.

### Overexpression *of hibris* is sufficient to disrupt R7 and R8 cell differentiation

To determine whether ectopic expression of *hbs* is sufficient to induce the expression of *Rh5* in R8 photoreceptor cells, we over-expressed *hbs* using the GAL4-UAS system [32] and the *P{GAL4-ninaE.GMR}* driver (FBtp0001315). **Fig 7A** shows that overexpression of *hbs* is sufficient to induce *Rh5* expression in many, but not all R8 photoreceptor cells. This occurs without perturbation of *Rh3/Rh4* R7 cell subtype ratio and is accompanied by mismatched *Rh4/Rh5* expressing R7/Rh8 photoreceptor cells pairs (not shown). To test whether the formation of these mismatched ommatidia could result from an inappropriate signal from *Rh4* expressing R7 cells or a defect in the default pathway and expression of *Rh6* in R8 cells, we overexpressed *hbs* in a *sev* mutant background that lacks R7 photoreceptor cells. **Fig 7B** shows that removal of R7 cells leads to a dramatic reduction but not elimination of *Rh5* expression. These results suggest that the ability of overexpressed *hbs* to induce *Rh5* expression in R8 cells is primarily or partially R7 photoreceptor cell independent.

### *hibris* is required non-cell autonomously for R7 and R8 cell differentiation

To determine whether *hbs* is required cell-autonomously in the R7 and/or R8 photoreceptor cells to enable normal paired expression of Rh3 and Rh5, we generated smaller *hbs*^*66*^ mutant clones in a heterozygous background as previously described [22]. Cells that are either wild type or heterozygous express a myristoylated, membrane associated GFP (*myr.GFP, P{GMR-myr.GFP},* FBtp0017435), whereas cells that are homozygous mutant for *hbs* do not express *myr.GFP* [33]. We dissociated ommatidia from animals constructed in this manner and counted the expression of *Rh5* versus *Rh6* in ommatidia that expressed *Rh3* in the R7 cell and in which the genotype of the R7 and R8 cells could be scored. **Fig 8A** shows a cluster of ommatidia from the experiment labeled with antibodies against *Rh3, Rh4, Rh5, Rh6*, with *myr.GFP* labeling shown in **Fig 8B**. Mispairing of *Rh3-Rh6* expression occurs in ∼ 20% of *cn*^*1*^ *bw*^*1*^ ommatidia that express *Rh3* in R7 cells (Fig 8C), consistent with previous results [11, 22, 23]. In the mosaic analysis, there is a statistically significant increase in the percentage of mispaired *Rh3-Rh6* expressing ommatidia ranging from 56 – 100%, regardless of the genotype of the R7 and R8 photoreceptor cells. Particularly noteworthy is the highly abnormal and pronounced effect on ommatidia in which both R7 and R8 photoreceptor cells are heterozygous or homozygous wild-type (R7+ R8+). These results unambiguously indicate that *hbs* is required non-cell autonomously for the establishment of paired opsin gene expression in the R7 and R8 photoreceptor cells. Although the severity of the mispairing is increased significantly in ommatidia in which R7 and R8 cells are both mutant (R7-R8-).

**Fig 8.**
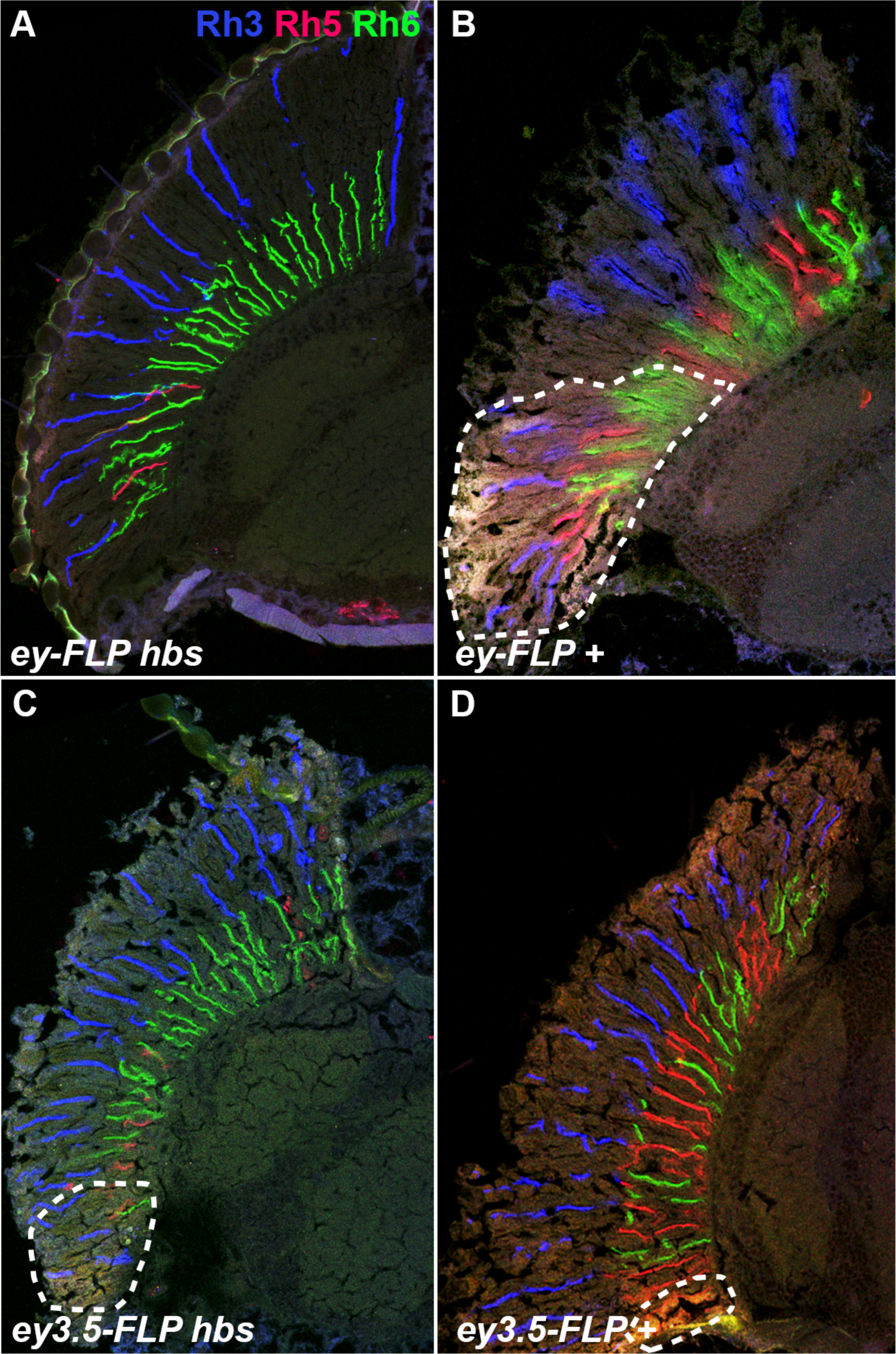
Opsin Expression in *hbs*^*66*^ Mutant and Wildtype Control Flies. Large FLP-FRT retinal clones were generated in the eye and optic lobes with *ey-FLP*, panels A and B, or in the retina alone with *ey3.5-FLP*. Homozygous *hbs*^*66*^ mutant clones are shown in panels A and C. Homozygous wild-type control clones (*+*) are shown in panels B and D. Heterozygous tissue is marked with *w*^*+*^ and outlined in panels B, C and D. *Rh3* (blue), *Rh5* (red) and *Rh6* (green) expression were detected by confocal microscopy with directly labeled monoclonal antibodies as described in **Materials and Methods**.

### *Notch* is required for R7 and R8 cell differentiation

*N* signaling plays an essential and reiterative role in the development of the compound eye [34], and is known to regulate *hbs* during myoblast fusion [35, 36], control cell adhesion in the eye through interactions with *hbs* [37], and *hbs* may also play a role in *N* cleavage [28]. To determine whether *N* is required for the establishment of paired opsin gene expression in the R7 and R8 photoreceptor cells, we reduced *N* activity at sequential stages of pupal development. We used shifts to a restrictive temperature of a temperature sensitive allele *N*^*l1N-ts1*^ (FBal0012887) in an otherwise *cn*^*1*^ *bw*^*1*^ background and compared with similarly treated *cn*^*1*^ *bw*^*1*^ controls. **Fig 9A** shows that at baseline without heat shock *N*^*l1N-ts1*^*; cn*^*1*^ *bw*^*1*^ flies have a significantly increased proportion of *Rh3:Rh4* expressing R7 cells compared to *cn*^*1*^ *bw*^*1*^ controls. The *Rh3:Rh4* ratio is significantly increased with heat shock at 24-36 hours after puparium formation (APF) and significantly decreased with heat shock at 36-48 hr APF compared to *N* mutants raised at the non-restrictive temperature. Because the variation of the *Rh3:Rh4* ratio in R7 cells would be expected to alter the *Rh5:Rh6* ratio in R8 photoreceptors, we specifically examined the percent of *Rh3-Rh6* mispairing as an index of impaired induction of *Rh5* expression. We found that *Rh3-Rh6* mispairing was significantly increased in both *N*^*l1N-ts1*^*; cn*^*1*^ *bw*^*1*^ animals and *cn*^*1*^ *bw*^*1*^ control animals heat shocked from 0-12 hr APF, **Fig 9B**. Statistically significant increases in *Rh3-Rh6* mispairing were noted at 24-36 hr APF, 36-48 hr APF and 48-60 hr APF in *N* mutant animals demonstrating a highly significant disruption in the paired expression of opsin genes in R7 and R8 photoreceptor cells. Because of heat shock induced lethality at 12-24 hr APF in *cn*^*1*^ *bw*^*1*^ control animals, results at this time point are difficult to interpret. However, these results conclusively demonstrate a requirement for *N* activity in regulating 1) *Rh3* versus *Rh4* expression in the pR7 and yR7 photoreceptors and 2) coupling of *Rh3/Rh5* expression in adjacent pR7 and pR8 photoreceptor cells of the same ommatidium.

**Fig 9.**
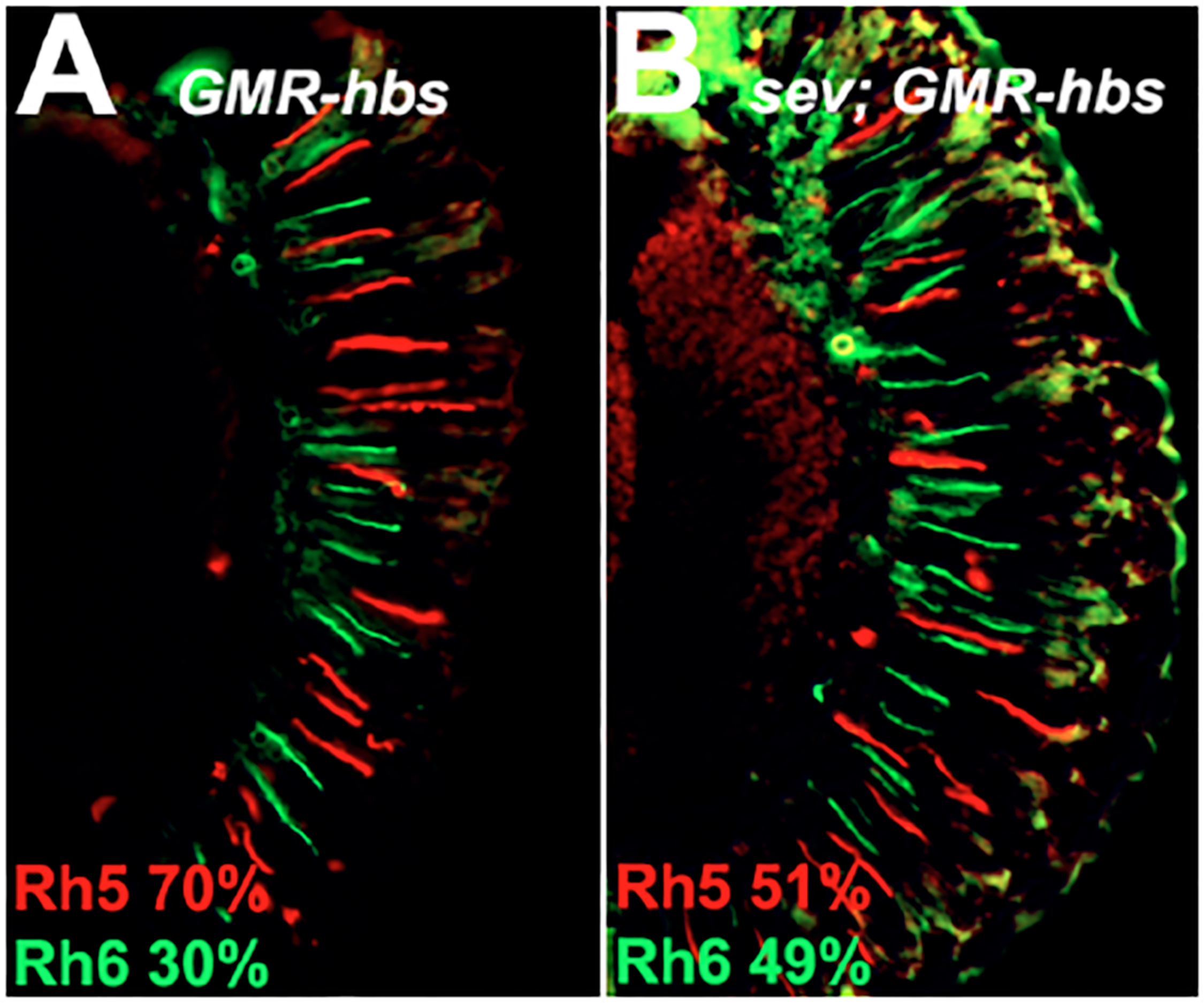
Overexpression of *hibris* Induces Increased *Rh5* Expression. Over expression of *UAS-hbs* with the *GMR-GAL4* driver leads to an increase in *Rh5* (red) expression, panel A. Removal of R7 photoreceptor cells (*sevenless*^*14*^ mutation) partially suppresses the effect, panel B. *Rh6* expression is shown in green.

## Materials and Methods

### Stocks and Genetics

Stocks were maintained in humidified incubators on cornmeal / molasses / agar media or standard cornmeal food with malt, and transferred on a rotating basis every three weeks as described [38-40]. *D. melanogaster* strains were obtained from individual laboratories or the Bloomington *Drosophila* Stock Center (BDSC). Genotypes were constructed using conventional genetic techniques, dominant markers and appropriate balancer chromosomes [39, 41].

### Genotypes of Animals Shown in Figures

**Figure 2 A, B, C:** *w*^*1118*^

**Figure 2 D, E, F**: *w*^*1118*^; *P{etau-lacZ}a69*

**Figure 5, Left column:** WT = *cn*^*1*^ *bw*^*1*^, **Right column:** *w*^*1118*^; *P{etau-lacZ}a69*

**Figure 6***: cn*^*1*^ *bw*^*1*^

**Figure 7***: y*^*d2*^ *w*^*1118*^ *P{ry*^*+t7.2*^*=ey-FLP.N}2 / w*^*1118*^*; P{w*^*+mW.hs*^*=FRT(w*^*hs*^*)}G13 L*^*\**^ */ P{w*^*+mW.hs*^*=FRT(w*^*hs*^*)}G13 hbs*^*113*0^

**Figure 8A:** *w*^*1118*^*/ y*^*d2*^ *w*^*1118*^ *P{ry*^*+t7.2*^*=ey-FLP.N}2 P{GMR-lacZ.C(38.1)}TPN1; P{ry*^*+t7.2*^*=neoFRT}42D hbs*^*66*^*/ P{ry*^*+t7.2*^*=neoFRT}42D P{w*^*+t\**^ *ry*^*+t\**^*=white-un1}47A l(2)cl-R11*^*1*^

**Figure 8B:** *w*^*1118*^*/ y*^*d2*^ *w*^*1118*^ *P{ry*^*+t7.2*^*=ey-FLP.N}2 P{GMR-lacZ.C(38.1)}TPN1; P{ry*^*+t7.2*^*=neoFRT}42D P{w*^*+t\**^ *ry*^*+t\**^*=white-un1}47A / P{ry*^*+t7.2*^*=neoFRT}42D P{w*^*+t\**^ *ry*^*+t\**^*=white-un1}47A l(2)cl-R11*^*1*^

**Figure 8C:** *w*^*1118*^*/P{w*^*+mC*^*=ey3.5-FLP.B}1, y*^*1*^ *w*^*\**^*; P{ry*^*+t7.2*^*=neoFRT}42D hbs*^*66*^*/ P{ry*^*+t7.2*^*=neoFRT}42D P{w*^*+t\**^ *ry*^*+t\**^*=white-un1}47A l(2)cl-R11*^*1*^

**Figure 8D:** *w*^*1118*^*/ P{w*^*+mC*^*=ey3.5-FLP.B}1, y*^*1*^ *w*^*\**^*; P{ry*^*+t7.2*^*=neoFRT}42D P{w*^*+t\**^ *ry*^*+t\**^*=white-un1}47A / P{ry*^*+t7.2*^*=neoFRT}42D P{w*^*+t\**^ *ry*^*+t\**^*=white-un1}47A l(2)cl-R11*^*1*^

**Figure 9A:** *w*^*1118*^*; P{GAL4-ninaE.GMR}12 / P{UAS-hbs.A*}

**Figure 9B:** *w*^*1118*^ *sev*^*14*^*; P{GAL4-ninaE.GMR}12 / P{UAS-hbs.A}*

**Figure 10:** *w*^*1118*^*/ y*^*d2*^ *w*^*1118*^ *P{ry*^*+t7.2*^*=ey-FLP.N}2 P{GMR-lacZ.C(38.1)}TPN1; P{ry*^*+t7.2*^*=neoFRT}42D hbs*^*66*^*/ P{ry*^*+t7.2*^*=neoFRT}42D P{w*^*+mC*^*=GMR-myr.GFP}2R*

**Fig 10.**
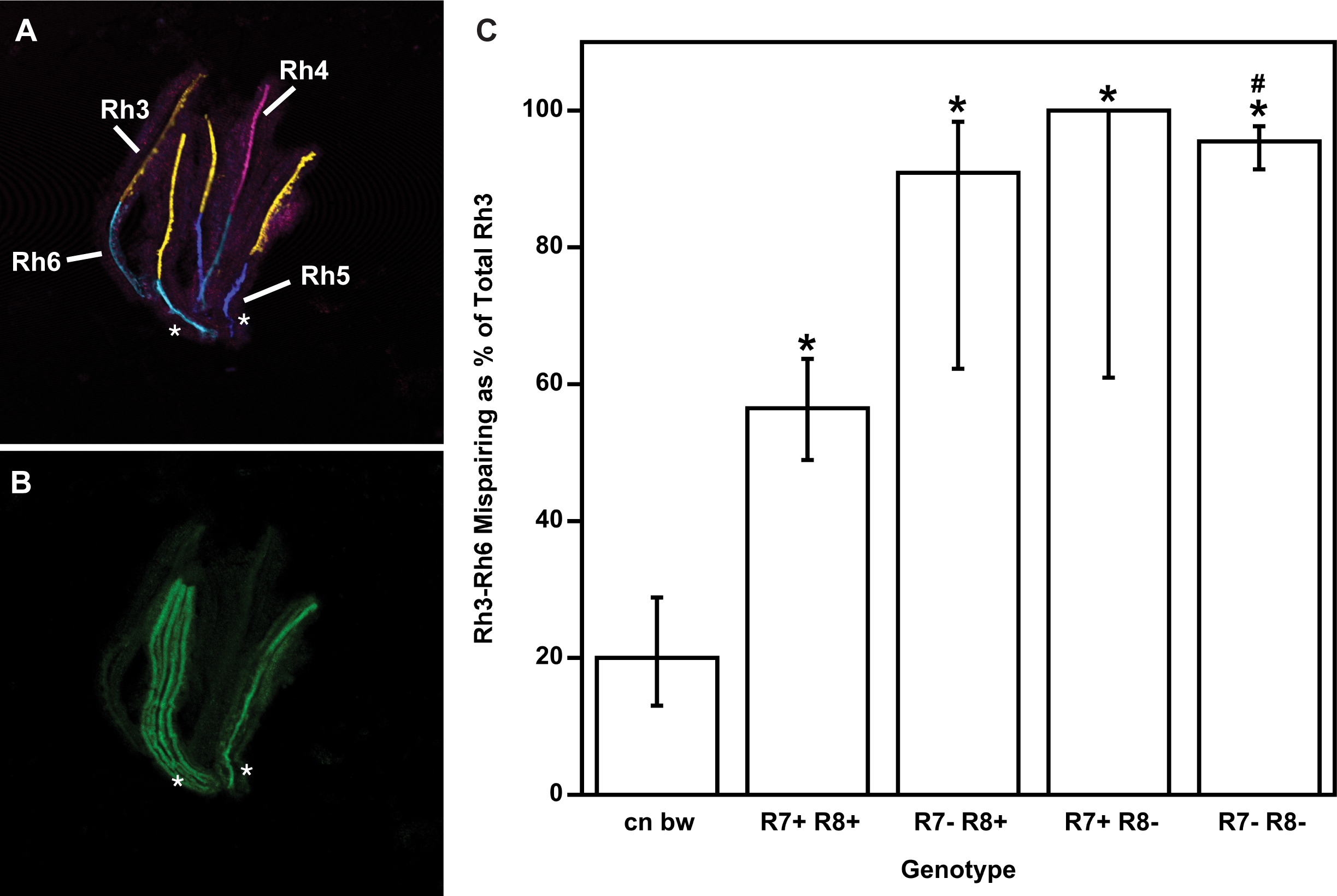
Testing for an R7 and/or R8 Cell Autonomous Requirement for *hibris*. Clusters of dissociated ommatidia from the eyes of *hbs*^*66*^ flies carrying small homozygous mutant clones were examined for expression of *Rh3* (yellow), *Rh4* (red), *Rh5* (blue), *Rh6* (magenta) panel A, and *myr.GFP* (green, panel B). Heterozygous or homozygous wild-type tissue is labeled with a myristoylated, membrane associated GFP (*myr.GFP*) (green, panel B, labeled R7+ or R8+ in panel C). Homozygous mutant tissue is unlabeled for *myr.GFP* (labeled R7- or R8- in panel C). Asterisks in panels A and B indicate two ommatidia showing *myr.GFP* expression in R7, R8 and other (in the case of the left ommatidia in panel B) photoreceptor cells. Panel C shows the percentage of mispairing of Rh3-Rh6 expressing R7-R8 cells in individual ommatidia (Y-axis) for unrecombined *cn bw* control flies, and each genotype of R7/R8 photoreceptor cells (X-axis). Asterisks (*) in panel C indicate that Rh3-Rh6 expression mispairing is significantly different statistically from the *cn bw* control (n=96 ommatidia) for R7+ R8+ (*p*=5.2×10^−9^, n=170 ommatidia), R7- R8+ (*p*=1.6×10^−6^, n=11), R7+ R8- (*p*=5.0×10^−5^, n=6), R7- R8- (*p*<10^−15^, n=178). Hashtag (#) in Panel C indicates mispairing is significantly different statistically from R7+ R8+ for R7- R8- (*p*<10^−15^, n=178). Error bars indicate the 95% confidence intervals for the measured percentages.

**Figure 11:** *cn*^*1*^ *bw*^*1*^ (*cn bw*) or *y*^*1*^ *N*^*l1N-ts1*^ *g*^*2*^ *f*^*1*^*; cn*^*1*^ *bw*^*1,*^ *(N; cn bw)*

**Fig 11.**
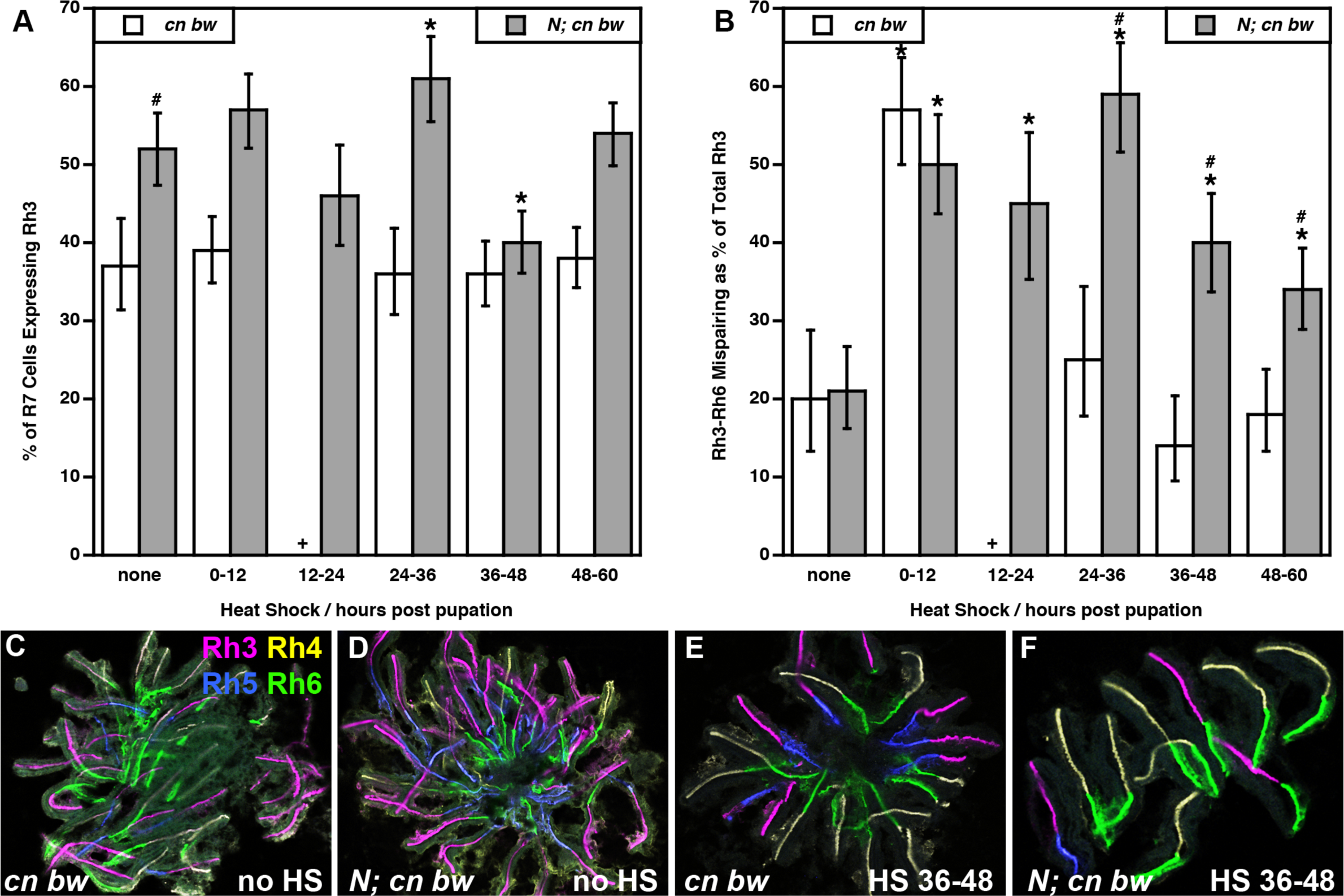
Analysis of *Notch* Function in R7 and R8 Photoreceptor Cell Differentiation. *N*^*l1N-ts1*^*; cn*^*1*^ bw^*1*^ (shaded columns) flies were compared to *cn*^*1*^ *bw*^*1*^ control (white columns) flies. Panel A shows data for the percent of R7 cells expressing Rh3 versus Rh4. Panel B shows data for the percent of Rh3-Rh6 mispairing in adjacent R7 and R8 photoreceptor cells within an individual ommatidium. Both panels A and B are bar graphs showing no temperature shift (none, raised continuously at 20°C) or a 12 hour temperature shift (temperature raised to 29°C at the indicated time in hours after puparium formation (APF) followed by return to 20°C). Hashtag (#) indicates significantly different from *cn bw* control receiving the same treatment *p*<0.05. Asterisk (*) indicates significantly different from the same genotype not heat shocked (none) *p*<0.05. + not recovered (heat shock induced lethality was observed for *cn bw* at this temperature shift time point). Error bars indicate the 95% confidence intervals for the measured percentages. Number of ommatidia counted for *cn bw* and *N; cn bw* respectively Panel A (no heat shock 259, 446; 0-12 hr heat shock 505, 411; 12-24 hr heat shock 0, 226; 24-36 hr heat shock 285, 306; 36-48 hr heat shock 509, 577; 48-60 hr heat shock 610, 586). Panel B (no heat shock 96, 232; 0-12 hr heat shock 197, 234; 12-24 hr heat shock 0, 104; 24-36 hr heat shock 103, 187; 36-48 hr heat shock 183, 231; 48-60 hr heat shock 232, 316).

### Immunohistochemistry

10μm cryosections were prepared and treated as previously described [11]. Dissociated ommatidia were prepared from six animals. Eyes were cut from heads using 28 gauge needles in Phosphate Buffered Saline (PBS). The retina, cornea +/- lamina tissue was shredded with needles, triturated 10 X with a 200 μL pipette tip and transferred to a microscope slide to dry at RT. Subsequent treatment was the same as cryosections. Antibodies were used at the following dilutions: directly conjugated mouse monoclonal anti-Rh5 (Texas Red, 1:100, RRID: AB_2736994) and directly conjugated mouse monoclonal anti-Rh6 (FITC, 1:100 RRID:AB_2736995) [42], mouse monoclonal anti-Rh4 (clone 11E6, 1:10, RRID:AB_2315271) [11, 42], mouse monoclonal anti-prospero (1:10, RRID:AB_528440, [43]), guinea pig polyclonal anti-senseless (1:1000, [44]), rabbit polyclonal anti-hibris (1:400, AS-14, RRID:AB_2568633, [45]). Secondary reagents were obtained from Life Technologies Corporation (Carlsbad, CA) or Jackson ImmunoResearch Laboratories, Inc. (West Grove, PA). An additional reagent was prepared from purified (Cell Culture Company, LLC, Minneapolis, MN) mouse monoclonal anti-Rh3 (RRID:AB_2315270). anti-Rh3 was directly conjugated using Alexa Fluor™ 647 Protein Labeling Kit (Invitrogen, catalogue number A20173) and used at 1:100 dilution. Immunofluorescence images were acquired with an Axioskop plus/AxioCamHRc (Carl Zeiss, Inc., Thornwood, NY) or by confocal microscopy using a Zeiss Pascal LSM (Carl Zeiss, Inc.) or Leica TCS SP5 (Leica Microsystems Inc., Buffalo Grove, IL).

### Statistical Analysis

Comparisons of the proportions (percentages) of opsin expression in different genetic backgrounds were performed with a z-score and are shown in **Table 1** and **Table 2** [46]. The *z*-score was calculated using the equation:

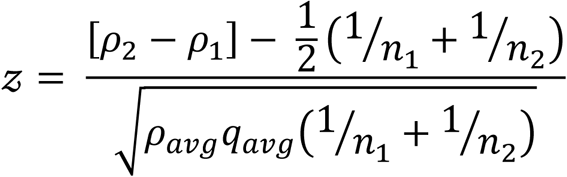

*p*_1_ and *p*_2_ = proportions of marker expression in each of the two different genotypes under comparison. *n*_1_ and *n*_2_ = number of ommatidia counted for each genotype. *pavg* = average proportion for both genotypes combined. *q*_*avg*_ = 1-*p*_*avg*_. The significance of the difference between the two proportions was determined from the normal distribution as a one- or two-tailed test. The 95% confidence interval of a proportion was calculated using the Wilson procedure without continuity correction [47, 48] using VasarStats [49].

### RNA *in situ* hybridization

Eye-antennal imaginal discs from third instar larvae were dissected in PBS, fixed in 50mM EGTA / 4% formaldehyde in PBS, rinsed in methanol, and stored in ethanol at -20°. Discs were treated with ethanol/xylene (1:1), rinsed with ethanol, post-fixed in 5% formaldehyde in PBS plus 0.1% Tween (PBT), washed with PBT, and digested with Proteinase K (5 μg/ml). Tissue was post-fixed again and pre-hybridized in hybridization buffer (50% deionized formamide, 5XSSC, 1 mg/ml glycogen, 100 μg/ml salmon sperm DNA, 0.1% Tween) at 48°C. Discs were hybridized overnight at 55°C with 2 μl digoxigenin-labeled antisense RNA probe in 100 μl hybridization buffer. Probes were prepared from cDNA clones D1 [50], GH09755 (FBcl0125531), GM02985 (FBcl014202), LD18146 (FBcl0156485), LP09461 (Fbcl0187603) of genes *hbs*, *pcs*, *CG10265*, *CG7639* and *ckn*, respectively. The hybridized imaginal discs were washed extensively with hybridization buffer at 55°C followed by PBT washes at room temperature. Discs were incubated with alkaline phosphatase-conjugated anti-digoxigenin antibody (1:2000, Roche Applied Science, Indianapolis, IN) overnight at 4°C. Discs were washed with PBT and gene expression was visualized with staining solution (100mM NaCl, 50 mM MgCl2, 100 mM Tris pH 9.5, 0.1% Tween) containing NBT/BCIP (Roche Applied Science). Stained imaginal discs were mounted and photographed using an Axioskop plus/AxioCamHRc (Carl Zeiss Inc.).

### *Notch* Temperature Sensitivity

Stocks of *y*^*1*^ *N*^*l1N-ts1*^ *g*^*2*^ *f*^*1*^ */ C(1)DX, y*^*1*^ *f*^*1*^*; cn*^*1*^ *bw*^*1*^ or control *cn*^*1*^ *bw*^*1*^ flies were maintained at 20°C. Male offspring were collected at the white prepuparium stage (P0). At 0, 12, 24, 36 and 48-hours after puparium formation (APF), pupae were shifted to 29°C for 12 hours and returned to 20°C until eclosion or formation of pharate adults. Retinas were dissected and ommatidia were dissociated and labeled with opsin antibodies for cell counting. 30 pupae were collected in each experimental or control group.

## Discussion

Here we describe the isolation and characterization of a novel allele of the *D. melanogaster* gene *hibris*, an evolutionarily conserved NPHS1 (nephrin) related IgSF member [51]. We show that *hibris* is required for the coordinated expression of opsin genes in adjacent R7 and R8 photoreceptor cells within the compound eye. Orthologues of this gene have been identified in many species, and numerous paralogues within species play diverse roles in organ system development and function [52]. Within the context of R7 and R8 photoreceptor cell differentiation and the regulation of opsin gene expression in the retinal mosaic, the specific functional role of *hbs* is unclear.

As noted briefly in the Introduction, the current model for the establishment of paired opsin gene expression in the R7 and R8 photoreceptors requires the type I activin receptor *baboon* (*babo,* FBgn0011300), bone morphogenetic protein type 1B receptor *thickveins* (*tkv*, FBgn0003716), transforming growth factor (TGF) beta type II receptor *punt* (*put*, FBgn0003169), many of their ligands, ligand processing convertases, and downstream effector enzymes [21]. In addition, the tumor suppressor kinase *warts* (*wts*, FBgn0011739), *hippo* kinase (*hpo*, FBgn0261456), *salvador* (*sav*, FBgn0053193), and *melted* (*melt*, FBgn0023001) a modulator of insulin/PI3K signaling [12], the *hpo* signaling cascade members *Merlin* (*Mer*, FBgn0086384), *kibra* (*kibra*, FBgn0262127), and the tumor suppressor *lethal (2) giant larvae* (*l(2)gl*, FBgn0002121) [19], and the transcription factors *ocelliless* (*oc*, FBgn0004102), *dorsal proventriculus* (*dve*, FBgn0020307) [53], *PvuII-PstI homology 13* (*Pph13*, FBgn0023489) [54] and *erect wing (ewg,* FBgn0005427) [55] are also required. Although not specifically tested in every case, all of these genes are thought to function cell autonomously within the R7 or R8 photoreceptor cells.

*hbs* is required in the eye for the induction of Rh5 expression based upon our experiments making homozygous mutant clones with *ey3.5-FLP* (**Fig 8**). However, in mosaic animals in which the R7 or R8 cells may be mutant or heterozygous in a mixed genotype environment, we find that *hbs* appears to have both cell autonomous and non-cell-autonomous effects. Specifically, ommatidia that carry R7 and R8 cells that are genotypically homozygous mutant are significantly more likely to show Rh3-Rh6 expression mispairing (**Fig 10C**, *p*<10^−15^). By contrast, in mosaic animals in which both the R7 and R8 cell of an individual ommatidium are heterozygous or homozygous wildtype, there is still a substantial reduction in Rh5 expression in R8 cells and a statistically significant (**Fig 10C**, *p*=5.2×10^−9^) increase in Rh3-Rh6 expression mispairing. This reflects a classic non-cell autonomous effect.

What could that effect be? Traditionally inductive processes are thought to occur between tissues or cells in which there is an inducer and a responder. Inductive signals are also often defined as instructive or permissive [56]. In the presence of an instructive interaction (i.e. from a pR7 cell), the responder (R8) develops in a certain way (as a pR8 cell expressing Rh5). By contrast, in the absence of the instructive interaction (yR7 or R7 cells absent, e.g. *seveneless* (*sev*) mutants), the responder (R8) does not develop in a certain way (does not become pR8 expressing Rh5, but rather becomes yR8 and expresses Rh6 instead as a default fate (with some exceptions [11]). If *hbs* played a formal instructive role in regulating the expression of Rh5 in R8 photoreceptor cells, then we would expect that its expression throughout the retina (*GMR-Gal4; UAS-hbs*) would lead to expression of Rh5 in all R8 photoreceptor cells even in the absence of R7 cells (Fig 9B). While the number of R8 cells expressing Rh5 is far higher than in *sev* mutants alone [10, 11, 22, 23], ectopic expression of *hbs* in this experiment is not sufficient to induce Rh5 expression in all R8 photoreceptor cells. Therefore, *hbs* does not play a strictly instructive role in this process.

As a potentially permissive regulator of R8 photoreceptor cell differentiation, *hbs* may play a role in establishing the architecture of the developing eye. Perhaps loss of *hbs* in mosaic or fully mutant animals disrupts cellular contacts that mediate signaling between R7 and R8. There is ample evidence for disruption of cone and pigment cell differentiation and eye roughening in *hbs* mutants [57, 58]. Furthermore, *hbs* and its binding partner *roughest* (*rst*) are known to have effects on axon guidance and synapse formation in the optic lobes [59-62]. Perhaps interactions within the lamina or medulla are responsible for some aspect of inductive signaling and expression of Rh5 in pR8. Finally, perhaps the loss of Rh5 expression in the *hbs* mutant eye reflects an inability to respond to the inductive signal, a loss of competence [63]. We previously suggested that *rhomboid* (*rho*, FBgn000463) and the *Epidermal growth factor receptor* (*Egfr*, FBgn0003731) may play a role in establishing competence of the R8 cell [22]. In these studies, we showed that *rho* is required for the induction of Rh5 expression in R8 photoreceptor cells, but like *hbs*, when *rho* is lost in mosaic retinas but the R7 and R8 cells of an individual ommatidia are wild type or heterozygous, there remains a dramatic effect on induction of Rh5 expression. Furthermore, loss of *Egfr* was also found to reduce the induction of Rh5 expression and also affect the proportion of pR7 and yR7 cells. These findings suggest that *hbs* likely plays a permissive, non-cell autonomous role in R7 and R8 differentiation.

Because *hbs* is both regulated by *N* signaling [50] and also thought to participate in *N* processing following its activation [28], our finding that *N* is also required for induction of Rh5 expression is not unexpected. This result is completely consistent with the previous findings that sequential loss of *N* signaling disrupted eye development and differentiation at every time point [34]. Similarly, alterations in *N* signaling also affect the proportion of pR7 and yR7 cells, suggesting that it as well as *Egfr* may regulate what is thought to be a cell-autonomous stochastic developmental process. These findings raise several notes of caution regarding the potential complexity of the system and the need to rigorously test the underlying hypotheses upon which the current model for R7 and R8 photoreceptor cell differentiation is based.

Subsequent analysis of the role of *hbs* in R7 and R8 photoreceptor cell differentiation will require further identification of its specific interaction partners in this system, either in the retina or optic lobes, as well as the temporal requirement for its involvement in R7 and R8 cell differentiation. Ample resources are available including available mutant strains [64], RNAi transgenics [65], and temporal and spatial mis-expression tools [66-70]. Despite these technical resources, defining the precise role of *hbs* in R7 and R8 differentiation will likely yield a complex system, reflecting coregulation of the IRM proteins [71], involvement of large complexes associated with scaffolding proteins [72], functional or genetic redundancy, compensation [73] and feedback.

## Acknowledgements

We thank Mary Baylies, Ruben Artero, Gerry Rubin, Amy Tang, James Mohler, and Jeff Sekelsky for *D. melanogaster* stocks, Mary Baylies for the *hbs* D1 cDNA clone, Karl Fischbach for the rabbit anti-hbs antibody (AS-14), and Hugo Bellen for the guinea pig anti-sens antibody. Stocks obtained from the Bloomington Drosophila Stock Center (NIH P40OD018537) were also used in this study. We thank Natalia Toledo Melendez for technical assistance, John Aldrich and Tom Jacobsen for comments on the manuscript and thoughtful discussion.

## Supporting Information

**S1 Table. Complementation of *a69* Recombinant Strains.** Recombinants described in Fig 3 were crossed to *a69* and the number of ommatidia counted expressing Rh5 or Rh6, Total counted, and % Rh5 are indicated in the table. Controls for comparison were homozygous *a69* mutants or *a69* / *w*^*1118*^ heterozygotes. Each recombinant strain was compared to both controls (right two columns) and was either not significantly different (NSD) or significantly different from (SDF) the indicated control at the *p* value stated. Statistical comparisons of strains were carried out as described in Materials and Methods. Controls are indicated at the bottom of the table. Recombinant strains having % Rh5 values intermediate between wild type and mutant phenotypes, but statistically significantly different from both, are shaded.

**S2 Table. Complementation of *a69* by Deficiency Strains.** A panel of thirty three deficiency stains were crossed to *a69* to test for complementation. The number of ommatidia counted expressing Rh5 or Rh6, Total counted, and % Rh5 are indicated in the table. The control for comparison was homozygous *a69* mutants. Compared to *a69* (right column) each deficiency over *a69* was either not significantly different (NSD) or significantly different from (SDF) *a69* at the *p* value stated. Statistical comparisons of strains were carried out as described in Materials and Methods. Values for the *a69* mutant are indicated at the bottom of the table. Deficiency strains failing to complement a69, which are not statistically significantly different from *a69*, are shaded.

**S3 Table. Complementation of *hibris* alleles by Deficiency Strains.** A panel of seven deficiencies were crossed to *a69*, *hbs*^*361*^, *hbs*^*459*^, *hbs*^*1130*^, *hbs*^*2593*^ and *cn*^*1*^ *bw*^*1*^ to test for complementation of the *a69* mutant phenotype. The number of ommatidia counted expressing *Rh5* or *Rh6*, Total counted, and % *Rh5* are indicated in the table. The control for comparison was homozygous *a69* mutants. The deficiencies failed to complement the tested genotype (shaded rows) or complemented the tested genotype (white rows). Complementation was defined as significantly greater *Rh5*% than (SGT) *a69* homozygous mutant at the *p* value shown using a one-tailed test. Statistical comparisons of strains were carried out as described in Materials and Methods. Values for the *a69* mutant are indicated at the bottom of the table. Crosses having results that differed from expected are noted (Exceptions).

**S4 Table. Strain Information.** Includes recombination stocks, deficiencies and alleles that complement a69. Stock genetics, Flybase ID and RRID are listed where available.

